# *In silico* assessment of arrhythmic risk following the implantation of engineered heart tissues in porcine hearts with varying infarct locations

**DOI:** 10.1101/2025.11.13.688201

**Authors:** Ricardo M. Rosales, Gonzalo R. Ríos-Muñoz, Ana María Sánchez de la Nava, María Eugenia Fernández-Santos, Javier Bermejo, Manuel Doblaré, Ana Mincholé, Esther Pueyo

**Affiliations:** Aragón Institute for Health Research (IISA), Zaragoza, Aragón, Spain; Aragón Institute of Engineering Research (I3A), University of Zaragoza, Zaragoza, Aragón, Spain; Bioengineering Department, Universidad Carlos III de Madrid, Leganés, Spain; Department of Cardiology, Instituto de Investigación Sanitaria Gregorio Marañón (IiSGM), Hospital General Universitario Gregorio Marañón, Madrid, Spain; Center for Biomedical Research in Cardiovascular Disease Network (CIBERCV), Madrid, Spain; Departamento de Medicina, Universidad Complutense, Madrid, Spain; CIBER-BBN, Instituto de Salud Carlos III, Madrid, Spain

## Abstract

Engineered heart tissues (EHTs) have shown promise in partially restoring ejection fraction after myocardial infarction (MI); however, their potential to introduce electrophysiological heterogeneities and promote arrhythmias remains underexplored. This study assessed the arrhythmogenic risk following immature EHT engraftment in infarcted ventricles using computational simulations that replicate preclinical protocols. EHT computational models were developed and integrated into nine validated porcine-specific biventricular models from pigs with left circumflex (LCx, n=4) or left anterior descending (LAD, n=5) MIs. Ventricular tachycardia (VT) susceptibility was evaluated using an S1-S2 stimulation protocol across varying pacing sites and coupling intervals, accounting for infarct characteristics, implantation site, conductivity, and the ventricular conduction system (CS). VT burden was quantified with a 0-1 inducibility score (IS). *In silico* reentrant activity reproduced the arrhythmic patterns observed experimentally in porcine MI models. VT vulnerability was greater in LAD than in LCx infarcts, consistent with a larger infarct size. Inclusion of the CS modified VT burden by providing conduction shortcuts that either facilitated or suppressed reentry. Remuscularization directly on the MI region (IS = 0.49) heightened VT inducibility in dense, transmural scars (IS = 0.16), whereas lateral EHT implantation (IS = 0.35) reduced this risk with respect to direct implantation. In non-transmural scars, VT inducibility varied with the implantation site. Matching EHT conductivity to host myocardium lowered or contained arrhythmogenicity (LCx-IS: from 0.5 to 0.25; LAD-IS: stable at 0.57). These results highlight the latent arrhythmic risk of EHT-mediated remuscularization after MI, identifying infarct substrate, EHT conductivity, and implantation site as critical determinants, and emphasize the importance of incorporating the CS for accurate risk assessment.

**Author summary:** Remuscularization with engineered heart tissue patches has shown preclinical potential to restore cardiac ejection fraction after myocardial infarction. However, the introduction of electrophysiological heterogeneities from these patches remains underexplored. To address this, we performed *in silico* investigations of post-infarction arrhythmogenesis and its progression after patch engraftment in porcine models with vessel-specific infarcts, applying a comprehensive arrhythmia inducibility protocol. We first computationally reproduced the experimentally observed arrhythmic patterns in pigs. Before patch engraftment, we found that larger infarcts increased ventricular tachycardia vulnerability. The inclusion of the ventricular conduction system modified arrhythmic outcomes, as electrical shortcuts could either facilitate or suppress reentry. Following remuscularization, highly transmural scars exhibited higher arrhythmicity, with lateral patch implantation mitigating it compared to direct placement. In contrast, for poorly transmural scars, the relationship between arrhythmia inducibility and patch location was unclear, highlighting digital twins as a tool for personalized risk prediction. When host-like patch conductivity was used, arrhythmia inducibility after engraftment was reduced or contained. Overall, our results: i) reveal the latent arrhythmogenicity associated with patch-mediated remuscularization of infarcted hearts; ii) identify infarction substrate, patch conductivity, and implantation site as key arrhythmogenic determinants; and iii) emphasize the importance of accurately modeling the cardiac conduction system.

## 1 Introduction

Cardiovascular diseases remain the leading cause of mortality worldwide, with ischemic heart disease (IHD) being the primary contributor, accounting for 33% of cardiovascular-related deaths in women and 40% in men annually [1]. In recent years, IHD mortality rates have increased in low- and middle-income countries, placing a significant economic burden through direct healthcare expenditures and substantial indirect costs, including productivity losses and unpaid care efforts [1]. This burden is projected to escalate due to the aging population of Europe and other regions, highlighting the urgent need for effective therapeutic strategies to treat IHD and its most prevalent manifestation, myocardial infarction (MI) [1].

MI occurs when prolonged ischemia, typically due to inadequate reperfusion, induces irreversible cardiomyocyte death and subsequent tissue injury. This loss of cardiomyocytes initiates a coordinated inflammatory response that ultimately replaces the necrotic myocardium with non-contractile fibrotic tissue, resulting in impaired systolic and diastolic function [2]. The structural remodeling associated with MI increases the susceptibility to heart failure and sudden cardiac death, likely linked to the development of sustained ventricular tachycardia (VT). The extent of post-MI cardiomyopathy is strongly influenced by the location of the infarct. MIs with occlusion of the left anterior descending (LAD) coronary artery generally lead to larger infarcts, more pronounced remodeling, and greater functional decline compared to MIs with occlusion of the left circumflex (LCx) coronary artery [3].

Current therapeutic options to prevent severe post-MI deterioration of cardiac function include heart transplantation and left ventricular assist devices [4, 5]. Although heart transplantation remains the definitive treatment for end-stage heart failure, its widespread application is hindered by a limited donor pool and significant post-operative challenges, such as primary graft dysfunction, immune rejection, and the need for lifelong immunosuppressive therapy [6]. Similarly, assist devices for the left ventricle (LV), commonly used as a bridge to transplantation, involve major surgical intervention and require prolonged anticoagulation, posing additional risks to patients [4, 5].

A promising tissue engineering strategy under active investigation is the remuscularization of the infarct scar using human-induced pluripotent stem cell-derived cardiomyocytes (hiPSC-CMs) to restore LV ejection fraction. Recent studies have shown that the implantation of engineered heart tissues (EHTs) containing hiPSC-CMs can improve the post-MI ejection fraction in athymic rats with LAD-induced infarctions [7]. However, hiPSC-CMs remain structurally and functionally immature compared to adult cardiomyocytes, even when cultured in biomimetic environments incorporating electro-mechanical stimulation and biochemical cues within bioprinted melt electrowriting scaffolds [7, 8]. Electrophysiologically, hiPSC-CMs exhibit spontaneous automaticity, elevated diastolic membrane potentials, prolonged action potentials (APs), and reduced upstroke velocities [9], resulting in suboptimal intercellular coupling and extended AP durations (APDs) in EHTs relative to the native myocardium [7, 8]. Consequently, EHT-based therapies can introduce additional electrophysiological heterogeneities, potentially increasing the risk of arrhythmogenesis following MI.

During the past few decades, advances in biological data acquisition, computational power, and data storage have propelled the emergence of *in silico* modeling and simulation as a powerful tool for investigating cardiac electrophysiology, particularly in the context of therapy evaluation and diagnostic support [10–12]. Computational electrophysiology offers significant advantages over clinical studies, including greater flexibility and substantially lower demands in terms of time, cost, and resources. To facilitate its adoption and scalability, simulations often rely on experimental data from animal models, which are more readily available than human datasets. In cardiac research, domestic *Sus scrofa* pigs are widely regarded as the gold standard due to their close physiological resemblance to humans, including similarities in coronary anatomy, hemodynamics, heart size, and electromechanical properties [13]. For example, detailed porcine computational models developed by Mendonca Costa et al. [14] and Gaur et al. [15] have demonstrated the utility of such models in supporting preclinical research while reducing reliance on animal experimentation. In this context, recent studies by Yu et al. [11] and Fassina et al. [16] have employed limited cohorts of human biventricular (BiV) and LV computational models, respectively, to characterize the electrophysiological impact and arrhythmogenic potential of post-MI remuscularization using EHTs.

This study aims to elucidate the mechanisms driving post-MI arrhythmogenesis and how they are influenced by EHT engraftment. To this end, we performed *in silico* investigations of chronic post-MI arrhythmogenesis and its progression after the engraftment of EHTs in multiple MI models with varying infarct locations. We used our nine previously developed, highly detailed porcine-specific MI models [17] and subjected them to a comprehensive arrhythmia inducibility protocol (AIP). First, we compared simulated arrhythmic dynamics with experimentally observed reentrant patterns obtained through optical mapping (OM). Next, we quantified VT burden across different infarct substrates, with and without the inclusion of the ventricular conduction system (CS), revealing key insights into the role of these factors in modulating arrhythmogenic potential. Furthermore, we performed a sensitivity analysis to determine how post-MI remuscularization with EHT influences arrhythmia susceptibility, thus identifying critical factors, such as EHT implantation site and electrical maturation, that can exacerbate or mitigate arrhythmic risk in this therapeutic context. Building on our previous work [18–20], we expanded our modeling framework by incorporating high-resolution porcine-specific BiV models and EHT geometries, the use of a novel pig-specific cellular electrophysiology model, and evaluations of the maturation state and site of implantation of the EHT. We also applied a comprehensive AIP in various MI morphologies and locations to robustly characterize arrhythmic risk.

## 2 Methods

Fig 1 shows the different steps of our proposed analysis. In summary, our nine previously developed MI personalized models were used in an evaluation of arrhythmogenesis before and after remuscularization therapy. In line with our previous work, we refer to the four LCx samples as pigs 4-7 and to the five LAD samples as pigs 8-12 [17].

**Fig 1.**
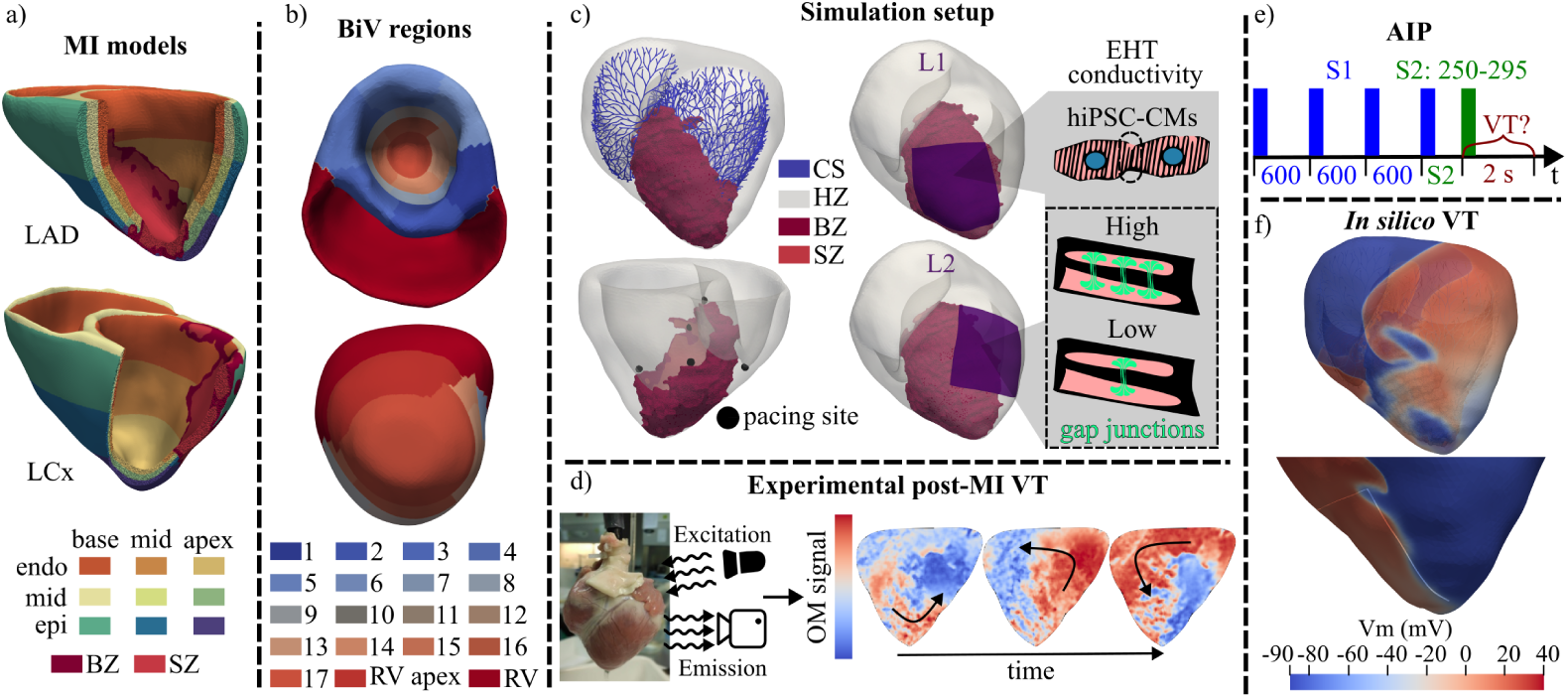
Summary of the arrhythmic evaluation performed before and after EHT implantation. a- Examples of BiV models of infarcts with LAD occlusion (top, pig 10) and LCx occlusion (bottom, pig 6). b- LV AHA segments and the apex of the right ventricle (RV) for another BiV model (pig 4). c- Example of a BiV model (pig 10) with its BZ, CS, pacing sites for the AIP and EHT implantation locations and maturation states. d- Experimental OM of the VT induced in pig 8. e- AIP depicting S1-S2 times. f- *In silico* VT induced in pig 10 before (top) and after (bottom) EHT remuscularization.

### 2.1 MI modeling

In a previous study, we built nine porcine-specific MI BiV models [17], two of which are shown in Fig 1a. Briefly, they consisted of four LCx and five LAD BiV models with infarcts that were delineated from late gadolinium-enhanced (LGE) cardiac magnetic resonance (CMR). LGE-CMRs were segmented using previously implemented automatic algorithms [21] with subsequent manual corrections. Based on the gadolinium washout and, therefore, the brightness of the LGE-CMR, porcine ventricles were divided into healthy (HZ), border (BZ), and scar (SZ) zones [17]. BZ represents the boundaries of MI, containing a mixture of viable tissue and collagen, while SZ defines the dense, non-conductive collagen remodeling site. BiV models were segmented in the apico-basal and transmural directions by solving a multi-diffusion problem [17, 22]. Standard fiber fields were generated with a rule-based model, as described by Bayer et al. [23]. CS was generated for each model using a fractal tree algorithm and was further projected onto the ventricular wall, as described in previous work [24, 25]. CS-activated ventricular nodes were located at a maximum distance of 0.5 mm from CS endpoints [17].

### 2.2 Arrhythmia inducibility

#### 2.2.1 Experimental Protocol

As can be observed in Fig 1d, induced arrhythmias in pigs 8-12 were experimentally measured by OM. The experimental preparation and the OM setup are described in detail in [17, 26]. Briefly, the heart was extracted from the animal and placed on a Langendorff circuit for dye and motion-blocking infusion. Only one induction event was mapped per pig. To induce VT, a stimulation catheter delivered a train of stimuli consisting of 12 initial pulses at a cycle length of 1000 ms, followed by 5 s of pacing with progressively decreasing cycle lengths from 300 to 150 ms until successful induction was achieved. The stimulation catheter was located in the LV for pig 8, in the left atrium for pigs 9-11, and in the right atrium for pig 12. The OM recordings were processed as previously reported [17, 18].

#### 2.2.2 Computational Protocol

Our implemented AIP was inspired by the work of Arevalo et al. [10]. As shown in Fig 1b, each BiV model was automatically divided into 19 regions using LGE-CMR-based segmentation of the LV and right ventricle (RV). The LV was segmented into 17 regions according to the American Heart Association (AHA) standard and consisted of 6 basal, 6 midventricular, 4 apical segments, and the apex, while the RV was divided into two regions: the apex and the remaining RV (Fig 1b). Based on this partition, the centroid of each segment was calculated, and its nearest endocardial node was identified. Pacing sites were defined as the set of nodes located within a 1 mm-radius sphere centered on each of these endocardial nodes. For each pacing site, an S1-S2 protocol was applied. Four S1 stimuli were applied every 600 ms, and one extrastimulus S2 was applied at 250, 265, 280, or 295 ms after the fourth S1 stimulus (Fig 1e). All stimuli had an amplitude of 80 µA*/*cm^2^ and a pulse width of 1 ms. Taking into account the high computational cost of running these simulations for all high-resolution BiV models under different conditions, the number of pacing sites was reduced by manually selecting at least 5 of the 19 pacing sites, with the criterion that they were well distributed around the MI (Fig 2). In the LCx and LAD models, the LV apex and RV apex were consistently used as one of the selected pacing sites, respectively.

**Fig 2.**
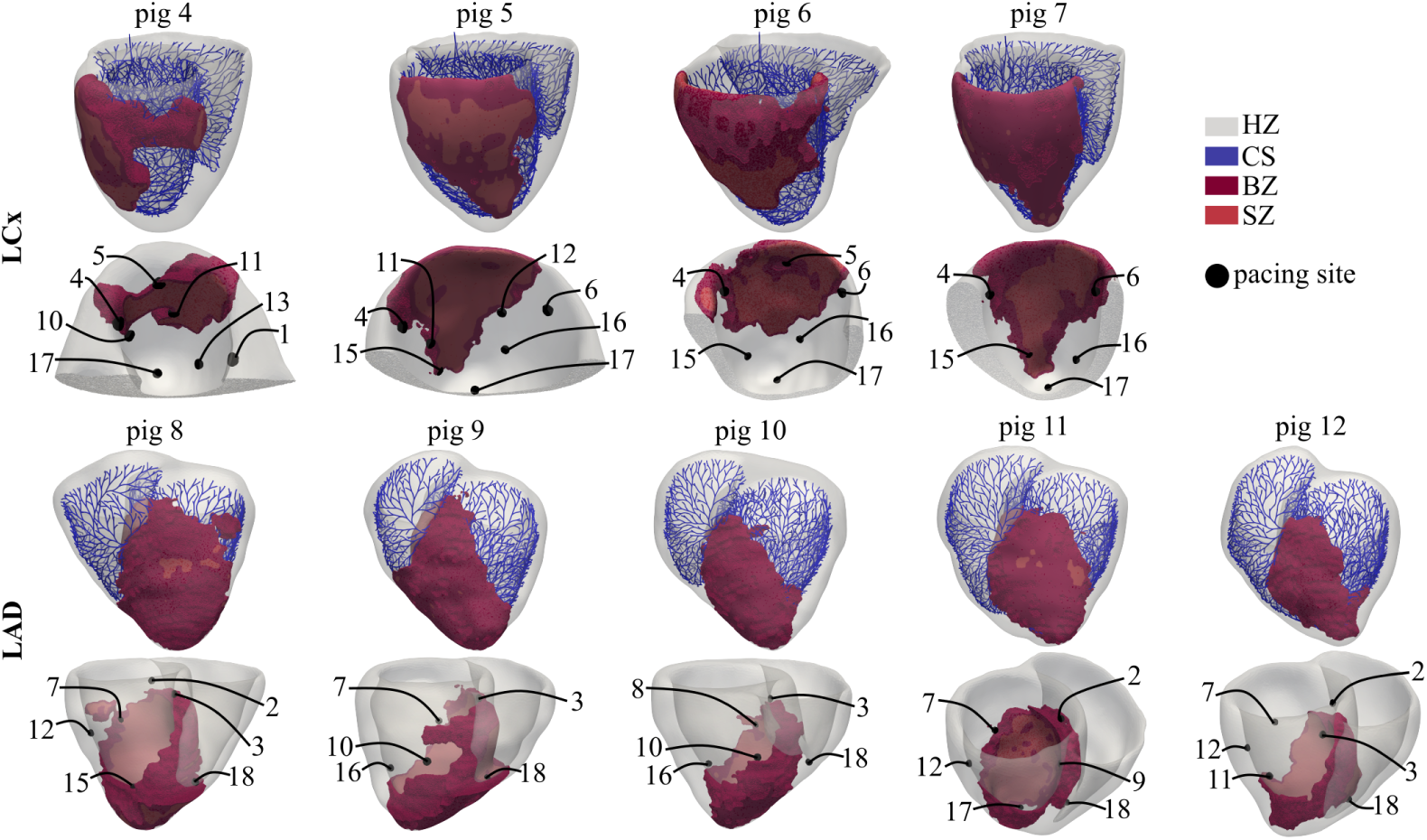
Porcine-specific BiV models of MI. BZ, SZ and CS are depicted together with the manually selected pacing sites used in each pig for application of the AIP. To show the pacing sites in LCx cases, a cut near the LV endocardial septum was performed and a sagittal-like view is presented.

The resulting electrical activity obtained from stimulating every pacing site was labeled as no reentry (NR), non-sustained VT (nsVT), and sustained VT (sVT). NR indicates cases where no reentrant activity was observed, while nsVT and sVT indicate that reentry vanished earlier than or was maintained for 2 s after the application of the S2 stimulus, respectively. From this categorization, an inducibility score (IS) was defined as 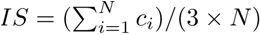, where *N* is the number of simulations used for its calculation, and *c_i_* is 0, 1, or 3 if the *i^th^* simulation resulted in NR, nsVT, or sVT, respectively. In this way, IS was defined in the range [0,1], with a value of 0 when the *N* simulations resulted in NR, 0.33 when all simulations resulted in nsVT, and 1 when sVT occurred in all cases.

### 2.3 Coupling of BiV and EHT models

A pipeline was implemented to build realistic BiV-EHT models using our previous work as a basis [19]. This pipeline is shown in Fig 3. First, the EHT was represented as a 40×40×1 mm^3^ squared superficial mesh, and a rigid transformation was used to bring it closer to the MI region of the BiV mesh (Fig 3a). The EHT size was selected based on experimentally achievable dimensions that allow remuscularization of most of the MI region [7, 27]. In the high-resolution EHT mesh, contact nodes were defined as those on the face closest to the epicardial surface. These contact nodes were individually projected onto the epicardial surface, with the same displacements applied to the nodes of the non-contacting face. The deformations imposed on the EHT, based on this node-wise contact displacement, resulted in an EHT deformation that followed the epicardial surface of the BiV mesh, as shown in Fig 3b. Subsequently, the EHT was embedded in the BiV model by projecting 0.5-0.6 mm of its thickness into the epicardium (see the right circle in Fig 3b). The projection direction was set as the inverse of the normal direction of the epicardial node closest to the centroid of the EHT mesh. Once embedded, the EHT mesh was merged with the BiV mesh using an *XOR* operation between the two triangular meshes (Fig 3c). The part of the EHT located inside the heart (blue region in Fig 3c) was removed, and irregular triangles that led to poor quality tetrahedralizations were cleaned with an isotropic explicit remeshing [28] of the BiV-EHT interface (Fig 3d).

**Fig 3.**
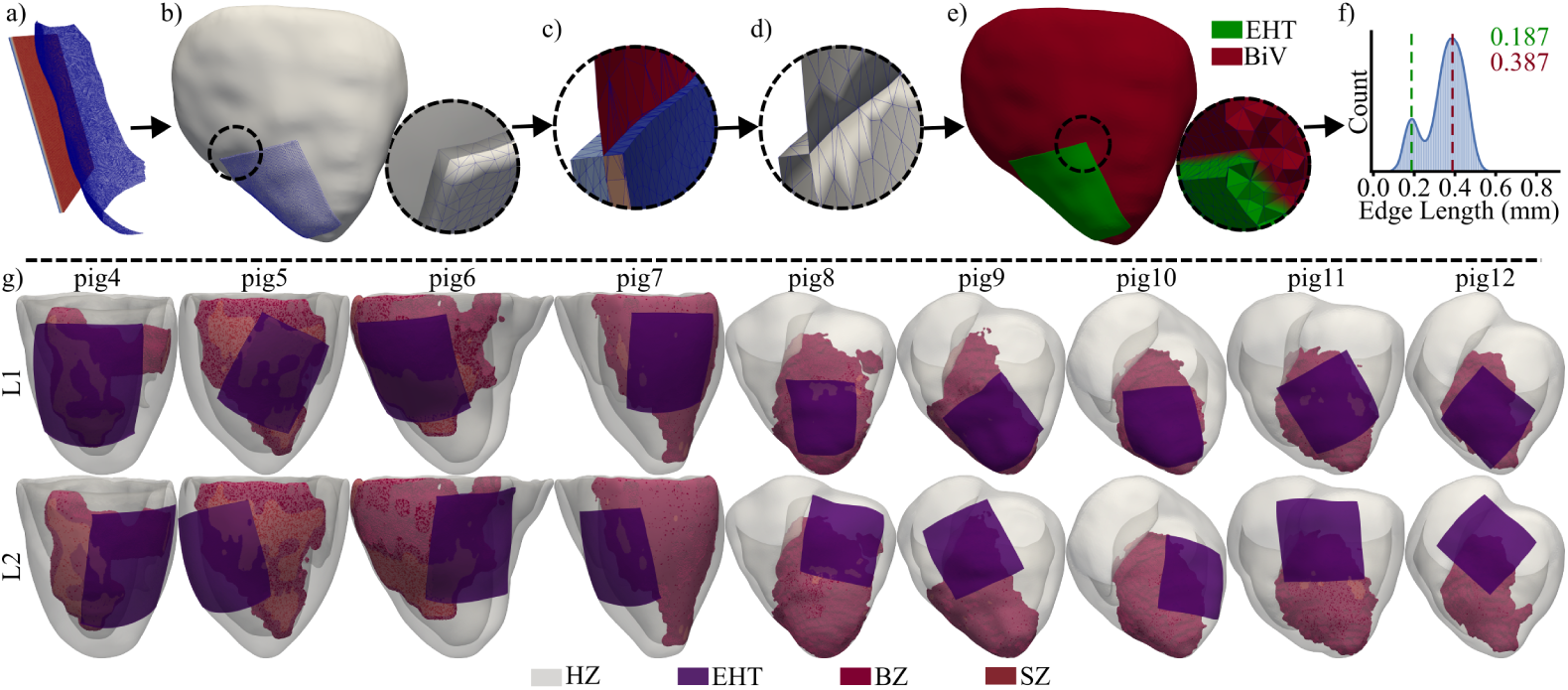
BiV-EHT modeling. Pipeline used to generate the BiV-EHT model for pig 10. a-Aligned EHT and epicardial meshes (EHT contact nodes in red). b- Deformed EHT mesh embedded on the BiV mesh (pig 10). c- Integrated BiV-EHT mesh with distinct regions and irregular triangles. Light-blue: outer EHT, blue: inner EHT, red: outer BiV, and pink: inner BiV. d- Cleaned BiV-EHT mesh. e- Tetrahedral mesh with BiV (red) and EHT (green) meshes. f-Bimodal edge length distribution in the BiV-EHT model. g- BiV-EHT models for all pigs and the two EHT locations.

The BiV-EHT meshes were discretized using Tetgen [29] (Fig 3e). The resulting tetrahedral meshes presented a bimodal edge length distribution, as shown in Fig 3f for pig 10. The low and high modes of these bimodal edge length distributions were calculated for all BiV-EHT models. A mean value of 187.2 µm was found for the low mode, which corresponded to the mean edge length in the EHT part. In addition, a mean value of 381.9 µm was found for the high mode, which corresponded to the mean edge length in the BiV part. The rationale for this domain discretization was to ensure simulation accuracy, particularly within the thin EHT and its surrounding regions. Due to its small thickness, the EHT required a higher element resolution, while the overall computational burden was kept to a minimum. The mean numbers of tetrahedrons and nodes for all models were 26 658 338*±*4 974 043 and 4 512 334*±*817 799, respectively. These values were reduced to 23 623 451*±*4 928 526 tetrahedrons and 4 054 310*±*815 100 nodes after SZ deletion to simulate this region as an insulator. The electrophysiological characteristics and fiber orientations of the BiV part in the coupled BiV-EHT models were interpolated from the corresponding BiV model. Random fiber orientations were assigned to the EHT based on prior evidence [11, 20].

### 2.4 BiV and EHT electrophysiology

The biophysically detailed cellular models of Gaur et al. [15] and Stewart et al. [30] were used to represent the AP of the porcine BiV models and their corresponding CS [17]. The electrical activity of hiPSC-CMs in the EHT was described by the updated cellular model by Paci et al. [31].

Regional repolarization heterogeneities in HZ were defined by tuning the conductance of the inward rectifier K^+^ current (*g_K_*_1_) in the Gaur et al. [15] cellular model, as this current is the main modulator of the APD in this model [15, 17]. In BZ of all models, *g_K_*_1_ was reduced to 29% of its default value to match the average prolongation of APD at 90% repolarization (APD_90_) reported after chronic MI [17].

All ventricular regions in the models were considered to have orthotropic conductivity with transverse isotropy. EHT conductivity was set to be completely isotropic due to the lack of precise information on connexin-43 distribution in hiPSC-CM cultures. Specifically, the longitudinal diffusion coefficients (LDCs) in BZ, HZ, and CS were set to 0.000882, 0.0013, and 0.013 cm^2^*/*ms (equivalent to 0.0882, 0.13, and 1.3 S*/*m for a membrane surface-to-volume ratio of 1000 cm*^−^*^1^ and capacitance per unit area of 1 µF*/*cm^2^), respectively [17]. The selected LDC value for BZ corresponded to the average of the individual values required to match *in silico* the conduction velocity in the MI region of each of the 5 LAD pigs measured by OM [17]. The excitability in BZ was reduced by decreasing the conductance of the fast Na^+^ current to 38% of its default value [10]. LDC for CS was reduced near the CS endpoints following a sigmoid curve to match the ventricular LDC, allowing physiological anterograde and retrograde electrical propagation between CS and ventricular tissue [32]. The transverse-to-longitudinal diffusion ratios were set to 0.345 and 0.25 in BZ and HZ, respectively. For BZ, this ratio reflects the reduced anisotropy observed relative to HZ in diffusion-weighted CMRs [17]. The ratio for HZ was based on measurements characterizing transverse anisotropy in healthy myocardium, where conduction velocity is greater along the longitudinal axis and lower along the transverse axis [33]. As transmembrane potential (*V_m_*) gradients have not been observed in experimental chronic MI electrocardiograms [17, 34], SZ was modeled as an insulator. This modeling approach avoids the *in silico* generation of *V_m_* gradients between the depolarized HZ and the electrically inactive SZ, which are not experimentally observed [17, 34].

All ventricular cell models were prepaced to steady state by applying periodic S1 stimuli (80 µA*/*cm^2^ magnitude, 1 ms duration, 600 ms period). For the Paci2020 model, prepacing was performed according to the stimulation described in [31] (0.55 nA magnitude, 5 ms duration) with a period of 600 ms. Stabilization of all cellular models was confirmed by verifying that APD_90_ remained constant for consecutive beats (CS: 310 ms under spontaneous automaticity, HZ: 196-218 ms, BZ: 384 ms, EHT: 405 ms).

### 2.5 Numerical simulations

First, simulations were conducted to investigate the contribution of MI characteristics and CS to arrhythmic risk. In the first group of simulations (G1), the complete AIP was applied to all BiV models. Specifically, S1-S2 stimulation was applied 4 times to each pig model in each selected pacing site, corresponding to the four S2 values of 250, 265, 280, and 295 ms. To validate our models, the simulated depolarization patterns calculated for LAD pig models were compared with those obtained from OM experiments for the same pigs when the experimental AIP described in Section 2.2.1 was applied. In this set of simulations, proarrhythmicity was evaluated as a function of the S2 delay, as well as the location, shape, and size of the MI.

In a second group of simulations (G2), proarrhythmicity was evaluated as a function of the inclusion or absence of CS in the models. Specifically, the AIP results after applying an S2 of 295 ms were compared for the LAD and LCx models with the highest inducibility (HI) and the lowest inducibility (LI), both with and without CS.

Subsequently, changes in vulnerability to arrhythmias after post-MI remuscularization with EHTs were investigated. A third group of simulations (G3) was carried out in which AIP was applied with an S2 of 295 ms before and after the engraftment of EHT in all pigs, and when the EHT was implanted in two different locations. For the central location L1, the EHT was implanted in the center of the MI. In contrast, for the lateral location L2, the EHT was positioned such that half of it overlapped the MI region, while the other half was in contact with HZ (Fig 3g). The rationale for selecting these locations was to balance two competing factors: mechanical support is believed to be maximized when the EHT is placed centrally within the MI; on the other hand, the survival of the EHT could be improved when it is in contact with well-vascularized healthy tissue. In all G3 simulations, the EHT conductivity was set to 10% (C10) of the value found in the HZ.

In addition, a fourth group of simulations (G4) was conducted to assess the effects of the electrical maturation of the EHT on proarrhythmicity. Higher maturation was represented by greater cell-to-cell coupling in the EHT. For G4 simulations, a pacing site was selected for each pig model based on the results of the G1 and G3 simulations; subsequently, S1-S2 stimulation was applied. Setting a value for the S2 stimulus of 295 ms at this pacing site, all coupled BiV-EHT models, with L1 and L2 locations for the EHT, were evaluated when the EHT conductivity was set to 90% (C90) of the value found in HZ. The results obtained for C10 and C90 were compared in each of the pigs.

In all simulations, the monodomain reaction-diffusion equation was numerically integrated using the open-source, in-house solver ELECTRA (v0.6.3) [35]. High-throughput execution was achieved through batch simulation using HERMES, the HPC Unit of the University of Zaragoza. Computed *V_m_* was saved at a time resolution of 1 ms, while the adaptive integration time step was constrained within the [0.01,0.1] ms range for all *in silico* calculations. In total, 335 simulations were performed. As some pacing sites did not capture any S2 stimulus, this involved the simulation of more than one million milliseconds of electrophysiological activity in high-resolution BiV models. Thus, this constitutes one of the largest and most complex cardiac electrophysiological simulation studies to date [10, 11, 14, 16, 36].

## 3 Results

### 3.1 Experimental and simulated arrhythmic patterns in MI

The experimental arrhythmic patterns mapped by OM in LAD pigs were compared with the results of the G1 simulations for pigs 8, 10, and 11 (Fig 4). In pigs 8 and 11, a counterclockwise reentry was observed in the anterior view of the OM recordings. These experimental patterns were reproduced by the *in silico* simulations when AIP was applied with an S2 of 295 ms at pacing site 12 for pig 8 and at pacing site 7 for pig 11. Generally, the depolarization wave traveled in the LV from the apex to the base, crossed to the basal RV, and propagated downward to the apical RV. In pig 10, the experimentally observed clockwise reentry was replicated *in silico* by stimulating pacing site 3 with an S2 of 280 ms. For this pig, the epicardial activation of the anterior view progressed sequentially from the LV apex to the RV base, then to the medial LV, and subsequently, the depolarization reached the laterobasal LV and the LV apex. Simulations also reproduced the counterclockwise reentrant activity observed in LAD pigs 9 and 12, with an S2 of 295 ms applied at pacing sites 16 and 12, respectively (see S1 Video).

**Fig 4.**
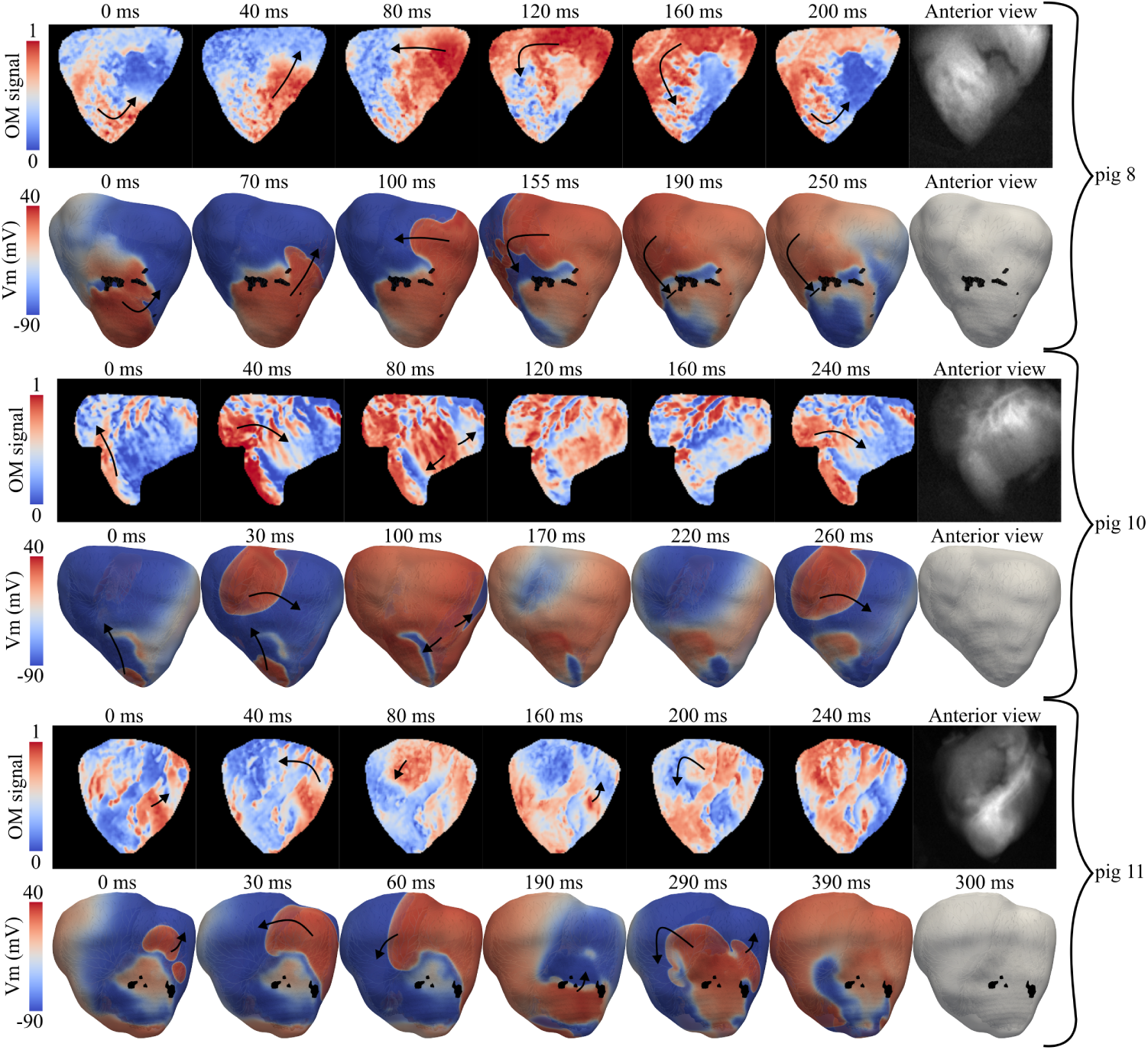
Reentrant activity in LAD pigs. VTs generated after application of the experimental (top) and *in silico* (bottom) AIPs.

Experimental VTs were sustained in all infarcted pigs with LAD occlusion, while nsVTs and sVTs were numerically observed for pigs 8-10 and 11-12, respectively. As an example, in pig 8, a single reentry was observed because the depolarization wave was stopped when traveling from the basal RV to the apical LV (see the simulated reentry of pig 8 in Fig 4). In spite of minor differences, experimental and simulated reentrant dynamics showed good agreement in all cases. Experimental and simulated arrhythmic patterns are provided for all pigs in S1 Video.

### 3.2 Quantification of VT inducibility after MI

Fig 5 summarizes the results of the G1 simulations, while the full set of results is provided in S1 Table. When AIP was applied, considering all pacing sites and all S2 values, pigs 8, 9, 10, 11, and 12 achieved IS values of 0.30, 0.37, 0.54, 0.7, and 0.1, respectively. Specifically, a single sVT was generated for pigs 8, 9, and 12, while 4 and 5 sVTs were found for pigs 10 and 11, respectively. Of all pigs with simulated LAD occlusion, pig 11 was the one with HI, while pig 12 was the one with LI. In particular, for pig 12, the application of AIP in 84.6% of the paced sites failed to develop a reentry. In addition, LCx pigs were found to be less vulnerable to arrhythmias. Only nsVTs were generated in pigs 5 and 7 (IS: 0.33), and NRs were observed in all cases in pigs 4 and 6 (IS: 0).

**Fig 5.**
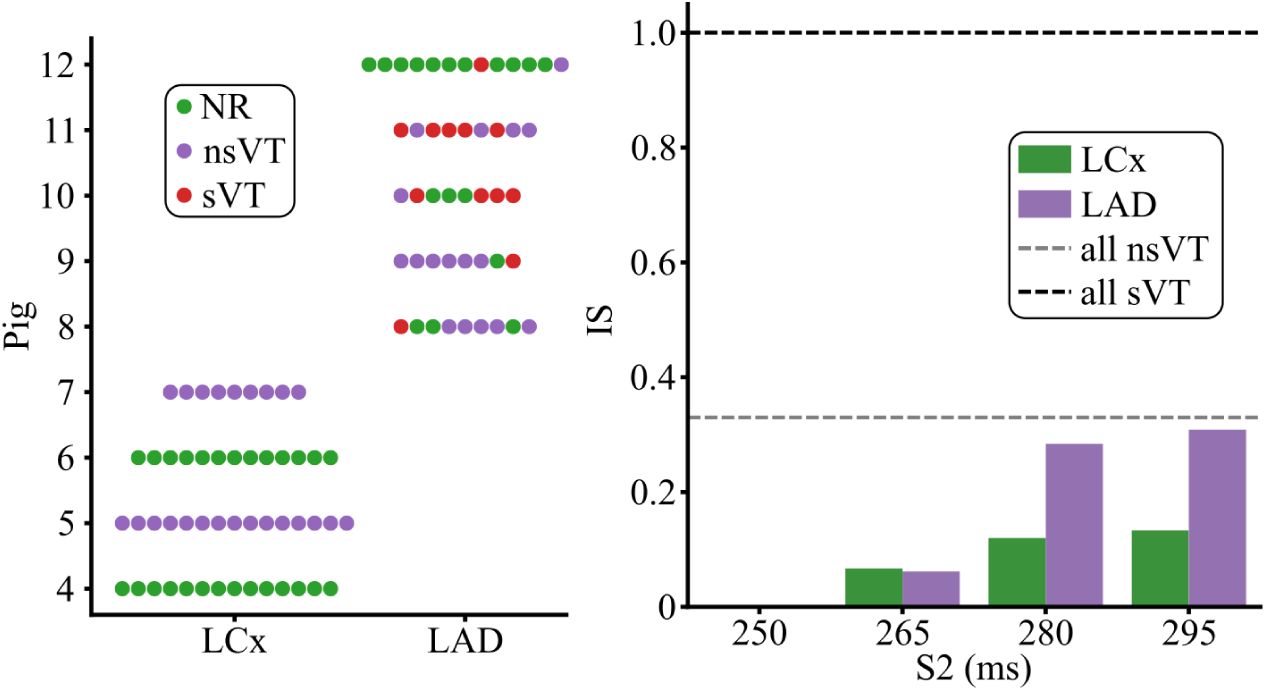
Post-MI inducibility. VT inducibility results as a function of MI location (left) and IS as a function of S2 values (right) for all 9 BiV models when all pacing sites were tested.

Regarding the S2 stimulus, the stimulated tissue was still refractory when the extrastimulus was applied 250 ms after the last S1 stimulus, preventing capture. Inducibility increased when the S2 delay increased, with contained increments in IS observed in LCx pigs. IS values of 0.06, 0.12, and 0.13 were measured for S2 values of 265, 280, and 295 ms, respectively, in LCx cases. In LAD pigs, IS increased sharply from 0.062 to 0.28 when S2 changed from 265 to 280 ms, and slightly from 0.28 to 0.31 when S2 changed from 280 to 295 ms, respectively. Thus, the S2 delay had a greater effect on inducibility in LAD pigs than in LCx pigs. For both types of MI, an S2 of 295 ms led to the highest inducibility. Therefore, other groups of simulations were conducted using this delay.

### 3.3 CS effect on arrhythmogenesis after MI

In the G2 simulations, LCx and LAD pigs with low and high inducibility in G1 were considered to evaluate the effect of CS on arrhythmogenesis. The complete set of results for the G2 simulations is provided in S2 Video and S2 Table.

For the group of LCx pigs, pig 6 exemplified a case of low inducibility in which CS played a fundamental role in blocking the reentry. As shown in the upper left panel of Fig 6, the application of AIP at the pacing site 6 generated an S2-derived depolarization wave that initially propagated from the anterior to the posterior LV through the apex. Then, a CS-derived epicardial breakthrough blocked this S2-derived activation in the laterobasal region, preventing reentry. Without the effect of the CS, the same S2-derived activation was not blocked by the epicardial breakthrough, allowing sVT. Pig 7 was chosen as an example of a high inducibility case (pig 5 was not chosen, as nsVT in pig 5 propagated through the low conducting laterobasal isthmus in the LV and could potentially vanish in a simulation of the entire heart, as described in the following Section 3.6). The application of AIP at pacing site 17 of pig 7 resulted in nsVT for a simulation with CS and in NR for a simulation without CS. In the first case, CS enabled the MI to function as a capacitor: the faster depolarization and subsequent repolarization of the MI region allowed slow loading of this region with S2-derived activation, followed by the release of the capacitive energy stored in the MI region from its apex back to HZ once the refractory-based impedance in HZ was low enough to allow the MI to be discharged, producing the reentry. Without CS, S2-derived activation did not penetrate the MI region, as it remained in its refractory period.

**Fig 6.**
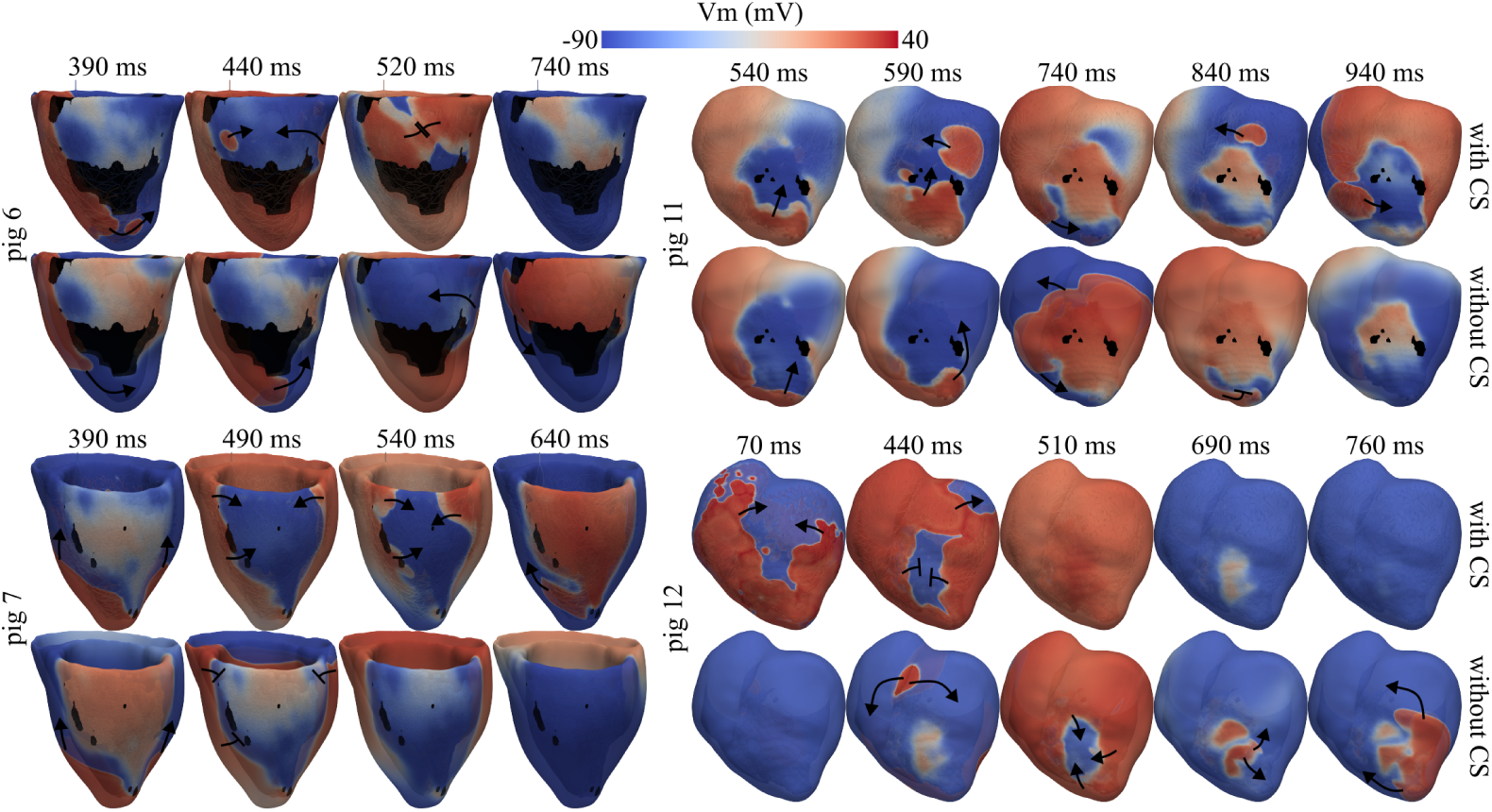
Changes in proarrhythmicity associated with CS. Results for pig 6 (LCx LI) stimulated at the pacing site 6, pig 7 (LCx HI) stimulated at the pacing site 17, pig 12 (LAD LI) stimulated at the pacing site 3 and pig 11 (LAD HI) stimulated at the pacing site 7 are depicted in the upper left, lower left, lower right and upper right panels, respectively. The timings refer to the time of application of the last S1 stimulus.

Similarly to the LCx cases, the presence or absence of CS determined an inversion in the occurrence of reentry in LAD pigs. In pig 11, associated with HI for LAD pigs, CS initiated delayed endocardial activations when pacing at site 7. These activations propagated transmurally, resulting in an epicardial breakthrough at 590 and 840 ms after the last S1 stimulus (see the top side of the upper right panel of Fig 6), sustaining VT. Beyond its role as an *electrical bypass*, CS also influenced VT dynamics through another mechanism linked to its intrinsic cellular properties. A nearly stalled and fading depolarization wavefront was reenergized in regions adjacent to the Purkinje-muscular junctions (PMJs). During the resting phase, CS had a diastolic potential higher than that of the surrounding myocardium, increasing the *V_m_* of the nearby ventricular PMJ nodes, as seen in Fig 7. Moreover, because of the longer APD of CS (Section 2.4), activated PMJs could further increase *V_m_*in these nodes, especially in HZ, bringing them closer to the excitation threshold. Together, these mechanisms may have facilitated local depolarization, thus supporting VT sustainability. In contrast, without CS, the re-energizing of fading depolarization wavefronts and, more importantly, the *electrical bypass*-derived epicardial breakthroughs were absent (see the bottom side of the upper right panel of Fig 6), and VT became non-sustained, terminating approximately 1 s after the last S1 stimulus. In pig 12 (LI for LAD pigs), the presence of CS generated an *electrical bypass* between the septal base and the apex of both ventricles when AIP was applied at the septal base (pacing site 3). This bypass revealed that CS enabled earlier depolarization of the MI region compared to the case without CS (see the lower right panel of Fig 6). Consequently, the S2 stimulus arrived at a completely repolarized MI 440 ms after the last S1 stimulus, resulting in NR. Without CS, a *capacitive effect* was observed, similar to that described for pig 7 with CS. The S2-derived depolarization wave partially halted in the MI and slowly penetrated this region. Later, reentry occurred when this MI-stored energy was unleashed back to HZ as this tissue regained excitability.

**Fig 7.**
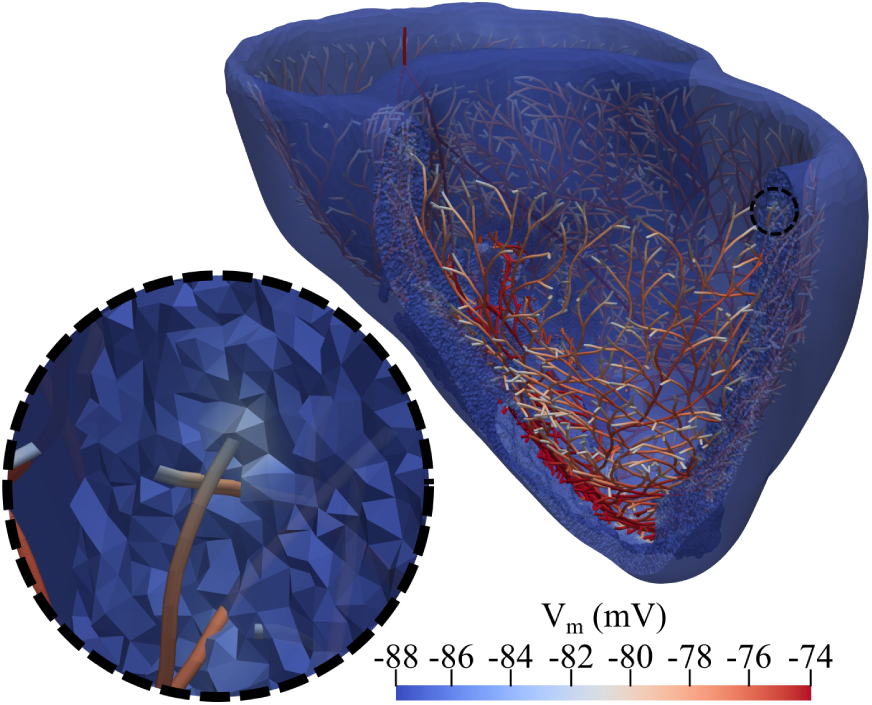
CS-BiV resting potential. Diastolic potential difference between the CS and BiV domain of pig 11, before applying the fourth S1 stimulus at pacing site 7.

In general, the presence or absence of CS drastically influenced arrhythmogenesis. As shown in Fig 8, removing CS decreased IS in the HI cases for both MI types (from 0.33 to 0.11 in LCx and from 0.66 to 0.33 in LAD pigs), but increased IS in LI cases (from 0 to 0.25 in LCx and from 0.22 to 0.44 in LAD pigs).

**Fig 8.**
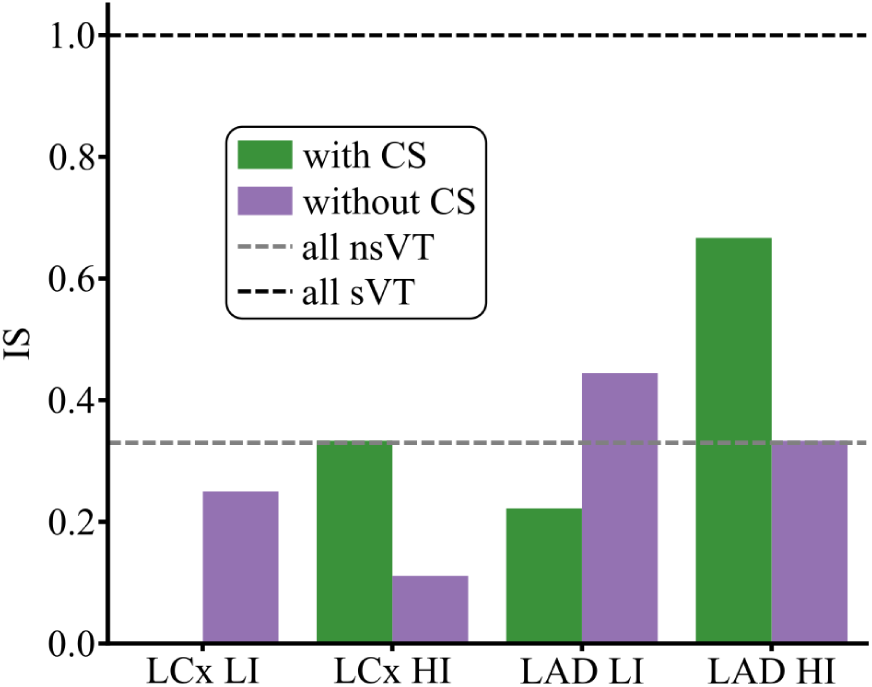
Effect of CS presence or absence on inducibility. IS computed with and without CS for LI and HI pigs of each MI type (LAD, LCx). LCx LI: pig 6, LCx HI: pig 7, LAD LI: pig 12 and LAD HI: pig 11.

### 3.4 Impact of EHT implantation location on arrhythmogenesis after MI

Fig 9 shows the effect of EHT engraftment location on arrhythmogenesis. IS values after MI and subsequent EHT implantation highlight a strong dependence of arrhythmogenesis on the occluded artery (see Fig 9a). In LCx pigs, EHT at L1 markedly increased IS from 0.16 to 0.49, while implantation at L2 resulted in a smaller rise (IS = 0.35). In LAD pigs, IS was slightly lower after implantation at L1 (IS = 0.33) compared with both pre-implantation and implantation at L2 (IS = 0.38), as can be seen in Fig 9a. S3 Table collects all the results for the G3 simulations.

**Fig 9.**
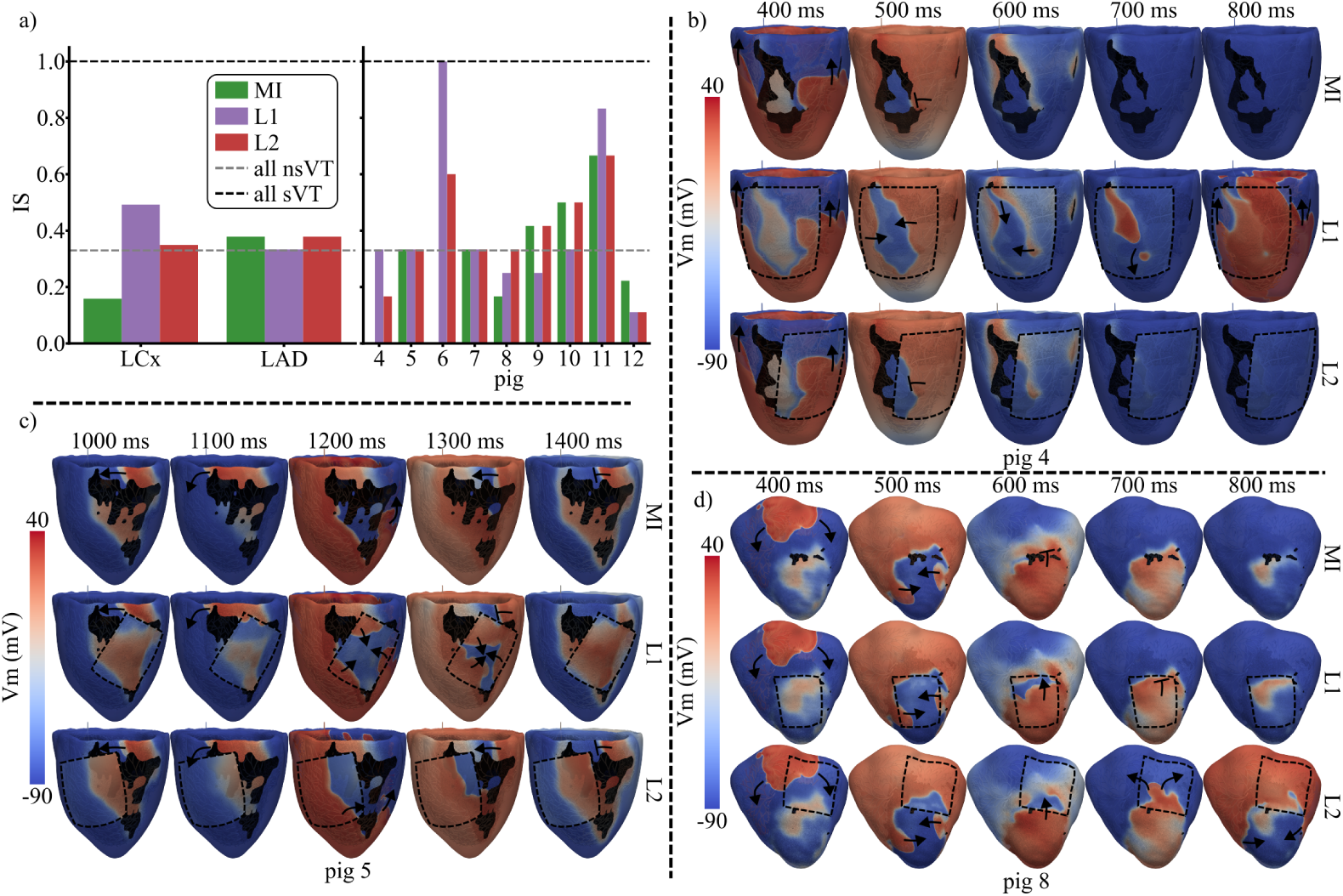
Effect of EHT engraftment location on arrhythmogenesis. In a, IS is presented for BiV MI models and for BiV-EHT MI models, with EHT located at L1 and L2. The left panel displays global results for LAD and LCx pigs, while the right panel shows results for each pig. In b, c and d, *V_m_* maps are presented for pigs 4, 5, and 8, with the EHT outlined (dashed black line). The depolarization-repolarization dynamics after AIP application are presented for MI (top), MI with EHT in L1 (middle) and MI with EHT in L2 (bottom). The pacing sites are 17 for pig 4, 6 for pig 5 and 3 for pig 8. The timings of the *V_m_* maps refer to the time of application of the last S1 stimulus.

Analyzing individual LCx pigs, IS remained stable in pigs 5 and 7 after MI and following EHT implantation at both L1 and L2. In pigs 4 and 6, EHT remuscularization increased IS, and its highest value was always found in the L1 position. For pigs 4, 5, and 6, the EHT covered the SZ extensively in the L1 position, which generated new conductive pathways between the basal and apical regions of the LV free epicardial wall. This phenomenon is illustrated for pig 4 in Fig 9b, for pig 5 in Fig 9c, and for pig 6 in Fig 10a.

**Fig 10.**
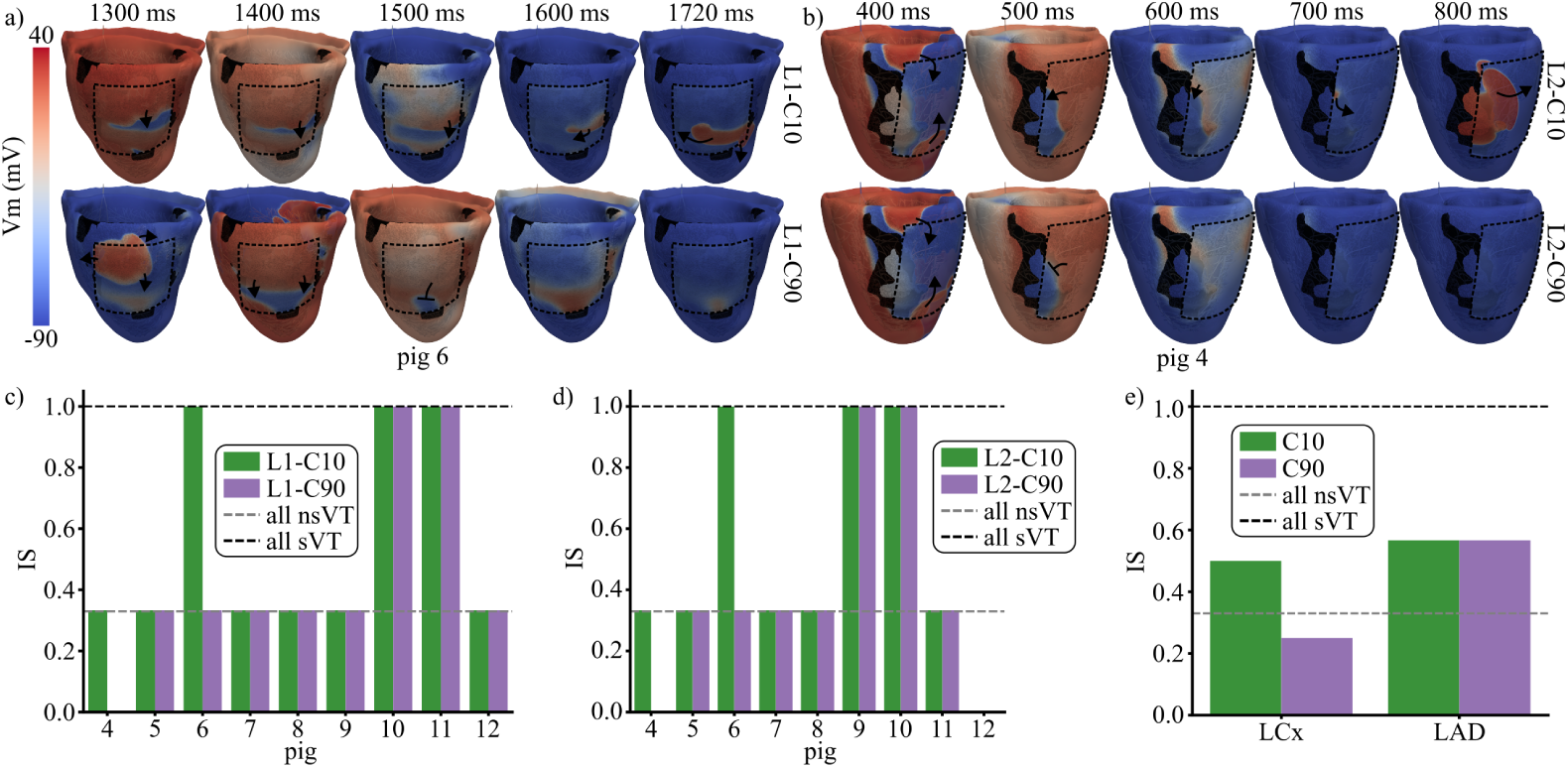
Effect of EHT electrical maturation on arrhythmogenesis. AIP-derived electrical activity is shown for pig 6 (a) and pig 4 (b) when low (top) and high (bottom) EHT (dashed black line) conductivities were used. The pacing sites were 15 for pig 6 and 1 for pig 4. The timings refer to the time of application of the last S1 stimulus. IS was obtained per pig using different EHT conductivities and locations L1 (c) and L2 (d). Panel e shows IS for conductivities C10 and C90 of the EHT in MI groups LCx and LAD.

Specifically, in pig 4, the S2 stimulus initiated at pacing site 17 propagated first in the apico-basal direction and then reached the central island of BZ, generating reentry when the EHT was located at L1 (middle row Fig 9b). In contrast, NR was observed for the same pacing site both after MI and with EHT at L2, as the S2-derived depolarizing wave at the latero-basal LV was unable to cross to the central BZ due to the lack of EHT-mediated electrical bridging. A similar phenomenon occurred in pig 6. When AIP was applied at pacing site 15, CS blocked reentry after MI, as shown in Section 3.3. A new reentrant pathway was generated in the base-to-apex direction when EHT was implanted at locations L1 and L2, as can be observed for L1 in the upper row of Fig 10a. This pig exhibited the largest IS increase after EHT engraftment. Across all five pacing sites assigned to this pig, AIP simulations yielded no reentry before remuscularization (MI: 5 NR) but resulted in reentry at all sites after remuscularization (EHT-L1: 5 sVTs, EHT-L2: 3 nsVTs and 2 sVTs). Electrical bridging by the EHT was also observed in pig 5 at the L1 location (see Fig 9c), although IS remained unchanged regardless of EHT presence or location (Fig 9a). This stable IS was probably dependent on the fact that this pig persistently presented a reentry blocking the depolarizing wave traveling from the base to the apex through the electrical bridge associated with the EHT, as illustrated in the middle row of Fig 9c.

Unlike LCx pigs, LAD pigs exhibited high variability in proarrhythmic responses following EHT implantation and EHT location (see Fig 9a). Interestingly, in LAD pigs, the implantation of the EHT in L1 reduced IS in 3 of the 5 pigs (pigs 8, 9, and 12) and increased it in the other two pigs. When the EHT was implanted in L2, IS was maintained in pigs 9-11, reduced in pig 12, and increased in pig 8. As an example, the depolarization-repolarization dynamics computed when AIP was applied at pacing site 3 in pig 8 are presented in Fig 9d. Here, similar electrical activity was observed after MI and after EHT implantation in L1, with S2-derived excitation penetrating the MI region and blocking in the mid-epicardial LV. In contrast, implantation in L2 led to reentry, as S2-derived excitation propagated into the highly excitable EHT and continued through the weakly depolarized wavefront, which had previously acted as a conduction barrier.

### 3.5 Contribution of EHT electrical maturation to arrhythmogenesis after MI

In the G4 simulations, EHT electrical maturation was modeled by a variation in its conductivity. As shown in Fig 10, increasing the EHT conductivity from C10 to C90 either decreased IS or kept it at baseline level.

Fig 10a illustrates this effect for two different pigs and EHT locations (pig 6, location L1; pig 4, location L2). In the case of pig 6 and location L1, increasing the conductivity of the EHT to C90 led to a homogeneous fast depolarization and subsequent repolarization of the EHT area above SZ, which prevented an EHT-mediated slow depolarization wave from maintaining the VT observed for the conductivity C10 of the EHT (Fig 10a, 1720 ms after the last S1 stimulus). A similar phenomenon was observed in the case of location L2 for the same pig 6, where the faster activation of the EHT transformed sVT into nsVT (see S4 Table).

For the case of pig 4 and location L2, increasing the conductivity of the EHT to C90 led to a smoother depolarization wavefront (Fig 10b, 600 ms after the last S1 stimulus). For the conductivity C10 of the EHT, a small self-standing activation was detached from the depolarizing wave and arrived at the peninsular BZ area, which maintained the VT, as a new depolarization wavefront was generated and amplified from this region (top Fig 10b). Increasing the conductivity of the EHT to C90 prevented any reentry from occurring in both EHT locations.

For the remaining LCx pigs (5 and 7), as well as for all LAD pigs, increasing EHT conductivity had no effect on IS compared to the low EHT conductivity setup, as can be observed in Fig 10c and Fig 10d.

When grouped by infarct type (LAD and LCx), higher EHT conductivity reduced the vulnerability to arrhythmias for LCx infarcts (IS from 0.5 for C10 to 0.25 for C90) but had no effect in LAD infarcts (IS of 0.57 for C10 and C90) (Fig 10e). S4 Video presents the complete electrical activity for pigs 4 and 6 after implanting the EHT in locations L2 and L1, respectively.

### 3.6 Role of MI characteristics on arrhythmogenesis before and after EHT implantation

The overall MI size, encompassing both SZ and BZ, was generally larger for LAD pigs than for LCx pigs, as can be observed in S5 Table. The ratio between MI and BiV volumes was 8.8%, 8.0%, 10.6%, and 14.0% for LCx pigs 4-7, and 30.2%, 27.1%, 16.6%, 20.8%, and 13.4% for LAD pigs 8-12, respectively.

For LAD pigs 8-11, a positive correlation between the volume of MI occupied by BZ and the value of IS was observed. Both IS (see Section 3.2) and the BZ/MI ratio increased progressively from pig 8 to pig 11 (BZ/MI: from 20.7% for pig 8 to 46.5% for pig 11). Pig 12, however, had the lowest IS value of all LAD pigs, despite having an intermediate BZ/MI volume ratio (36.2%). For LCx pigs, pig 7 showed both the highest IS value and the highest BZ/MI volume ratio (45.5%). Pig 5 had the same IS value as pig 7 (the highest among LCx pigs); however, it presented a low BZ/MI volume ratio of 20.8%. This could be explained by the fact that the reentry observed in pig 5 spread along a slow-conducting isthmus located in the latero-basal region of the LV (Fig 9c), suggesting that it could disappear in a simulation of the entire heart in which the addition of the outflow tracts and the atrial tissue alters the isthmus.

In the context of MI remuscularization with EHT, we observed that the size of the epicardial SZ correlated with the magnitude of IS induced by EHT implantation. For LCx pigs, the epicardial SZ was higher in pigs 4-6 than in the other pigs (see Fig 2, Fig 3g, and S5 Table). The ratio between the epicardial area of the SZ and the entire MI was 48.6%, 42.9%, 47.4%, and 3.8% for LCx pigs 4-7, respectively. Correspondingly, pigs 4 and 6 experienced the greatest increases in arrhythmia vulnerability after EHT implantation, which were reduced by placing the EHT in location L2 and increasing the conductivity of the EHT in either of the two locations. Pig 5, despite a high epicardial SZ/MI ratio, showed no IS changes. In this case, the EHT created a new reentrant pathway from the base to the apex. However, unlike pigs 4 and 6, this EHT-derived reentry was blocked by the reentry that occurred before the engraftment of the EHT (Fig 9c). In LAD pigs, epicardial SZ/MI ratios were generally low, ranging from 0% (pigs 10 and 12) to 15.1% for pig 8.

## 4 Discussion

### 4.1 *In silico* models reproduce experimental porcine-specific arrhythmic dynamics

Our *in silico* simulations reproduced the experimental dynamics observed in OM recordings of Langendorff-perfused LAD porcine hearts, which consistently exhibited VT induction, predominantly sVT. This agreement supports the robustness and reliability of the proposed modeling pipeline. In simulations, nsVTs were observed in pigs 8–10, whereas sVTs were reproduced in pigs 11–12, aligning with experimental observations where sVT was always induced.

The reduced reentry speed observed in simulations compared to experimental rotors (Fig 4 and S1 Video) likely reflects the use of an average BZ conductivity across models, which could have attenuated local conduction. This averaged conductivity was calibrated against conduction velocities in the five available LAD pigs and subsequently generalized to all LAD and LCx models. Personalization was not feasible for LCx pigs due to a lack of experimental data [17]. Similarly, CS distributions were assigned based on spatial benchmarks but lacked sufficient experimental validation [17]. Despite these limitations, the AIP simulations qualitatively captured the experimental arrhythmic dynamics, supporting the robustness of the modeling framework.

### 4.2 An *in silico* protocol for post-MI arrhythmia vulnerability testing

Arrhythmia inducibility was assessed with an *in silico* AIP inspired by prior approaches, such as the Virtual Heart Arrhythmia Risk Predictor [10], but streamlined with fewer pacing sites, reduced S1 trains, and a single S2 stimulus. Since BZ is the key substrate for reentry [14, 37, 38], pacing sites were placed homogeneously around the MI border, as distant pacing tends to elicit similar reentries. Under deterministic modeling, electrophysiological activity stabilized after a few beats, justifying the shorter pacing protocol. With this strategy, the simulations produced extensive reentry patterns closely resembling the experimental ones (see Section 3.1).

S2 intervals ranged from 250 to 300 ms, consistent with earlier studies [10, 14, 39, 40]. However, due to HZ/BZ refractoriness, as defined by the Gaur et al. [15] cellular model and its experimentally derived MI phenotype, short S2 intervals or near-MI pacing often failed to capture, preventing VT induction. Longer S2 delays increased inducibility, and variability in such inducibility across S2 values highlighted the value of selecting the shortest effective S2 and identifying the S2 delay that produces the highest inducibility for post-MI remuscularization analyses (G3 and G4 simulations). This establishes a robust framework for assessing inducibility with the Gaur et al. [15] model embedded in BiV simulations.

LAD pigs were consistently more arrhythmogenic than LCx pigs, with each presenting at least one sVT. This finding aligns with clinical observations of a greater incidence of arrhythmia in LAD infarction compared to LCx [41], likely reflecting the larger infarct size and remodeling observed in LAD cases [3]. Our simulations further showed that a larger BZ extent within the infarcted region was associated with greater arrhythmia vulnerability, consistent with prior canine and human studies [37, 38]. Exceptions likely arise from BiV modeling constraints, which may introduce artificial low-conduction pathways that sustain reentry (e.g., pig 5) or CS architectures that effectively block reentry (e.g., pig 6).

To our knowledge, this is the first computational study to reproduce the experimentally higher VT burden associated with LAD infarction using biophysically detailed and vessel-specific porcine MI models.

### 4.3 CS as a modulator of post-MI arrhythmia susceptibility

This study identifies the CS as a key determinant of post-MI arrhythmia susceptibility. Although the CS has been recognized as a potential modulator of conduction [42, 43], its specific role in post-MI proarrhythmicity remains unexplored. Using anatomically realistic CS distributions [17, 42], we demonstrate that CS can strongly influence arrhythmic dynamics by creating *electrical bypasses* across ventricular regions. These electrical shortcuts may sustain or terminate reentry or alter MI depolarization timing via the described *capacitive effect*.

Our findings align with clinical evidence reporting CS-induced VTs in ischemia, including reentrant bundle branch VT, involving the entire ventricular CS [44], and Purkinje-based intramyocardial VT, primarily involving the left posterior Purkinje fibers [44, 45]. Regarding the latter, Bogun et al. [45] identified a plausible Purkinje involvement in post-MI sustained monomorphic VT based on relatively narrow QRS complexes with left or right bundle branch block morphology that meet the following criteria: concealed entrainment at sites exhibiting Purkinje potentials; VT cycle length variability following changes in intervals between Purkinje potentials; or reproduction of the VT morphology through pacing at a site showing Purkinje potentials during sinus rhythm. Garcia-Bustos et al. [42] further confirmed the anatomical substrate for post-MI CS-derived VTs by showing that PMJs are capable of surviving reperfusion-related ischemia in chronic MI pigs.

Computationally, Berenfeld et al. [46] showed that reentries through Purkinje fibers can generate endo-epicardial breakthroughs that sustain polymorphic VT in a non-ischemic canine CS-BiV model, in line with our observations here. More recently, Sayers et al. [47] showed that MI-induced downregulation of fast-conducting connexins and alterations in ion channel expression in non-regenerative CS of postnatal mice slowed conduction and increased the incidence of arrhythmias in a BiV human model with LAD MI. Our study extends and complements these insights by testing CS effects across multiple ischemic substrates and pacing sites, showing how CS critically shapes reentry inducibility and dynamics.

Accurate CS modeling is, therefore, essential to prevent underestimation or overestimation of arrhythmic risk and to ensure the reliability of next-generation *in silico* arrhythmia assessments.

### 4.4 EHT conductivity has a major impact on arrhythmia vulnerability after MI remuscularization

Our simulations demonstrate that the conductivity of the EHT is a key determinant of arrhythmia susceptibility following MI remuscularization. Across the nine porcine-specific BiV models and the two implantation sites, greater and more biomimetic intercellular coupling consistently reduced or prevented increases in arrhythmia vulnerability. These results are consistent with previous computational studies using realistic ventricular representations [11, 16, 19, 20] and models of transmural MI slabs [18, 48], where higher EHT conductivity decreased VT burden and repolarization time gradients (a known proarrhythmic marker).

Specifically, Yu et al. [11] similarly reported a reduced VT burden in BiV-MI models when EHT electrical properties approached those of the host ventricular tissue. Fassina et al. [16] found that sVTs disappeared when conductivity reached nearly healthy levels, even in immature repolarization phenotypes. Likewise, our earlier studies showed reduced repolarization gradients with increasing EHT conductivity, irrespective of hiPSC-CM alignment or the degree of graft attachment [19, 20].

Taken together, these findings highlight that a mature EHT with high conductivity consistently mitigates, or at least contains, remuscularization-induced VT burden across diverse conditions involving cell alignment, degrees of graft attachment, infarct substrates, and implantation sites.

### 4.5 MI substrate strongly modulates arrhythmia risk after EHT implantation

Following post-MI remuscularization, arrhythmia inducibility exhibited marked artery occlusion dependence, varying substantially in LAD pigs but remaining more stable in LCx pigs. In LAD pigs, the average IS after EHT implantation resembled pre-engraftment values across both implantation sites. In contrast, LCx pigs showed a clear increase in average IS after EHT implantation, although lateral placement reduced reentry compared to central placement. These results extend prior observations by Yu et al. [11], who demonstrated location-specific VT incidence in two MI models where infarcts were confined to the endocardium or were highly diffuse in the epicardium, consistent with our findings in LAD pigs.

In pigs with highly transmural SZ, EHTs created novel isthmuses with fetal-like electrophysiology that, in some pigs (e.g., 4-6), enabled base-to-apex conduction and promoted unidirectional block and reentry. These proarrhythmic effects diminished with increased EHT conductivity, in agreement with previous studies [49]. In some cases, preexisting reentry blocked the EHT conduction path, invalidating any conductivity effects. In pigs where the EHT primarily covered conducting tissue (e.g., 7-12), no additional unidirectional blocks formed, as arrhythmicity was unaffected by conductivity changes. This suggests that inducibility after remuscularization in pigs without transmural SZ depends mainly on EHT-MI spatial relationships and the immature EHT phenotype.

Consistent with Campos et al. [49], we find that low conducting isthmuses are important for the onset of reentry, while reduced conductivity is the main contributor to its sustainability. This mechanism explains why post-remuscularization VT burden in weakly transmural infarcts varies strongly with EHT location at low EHT conductivity but is reduced or contained when EHT conductivity is close to biomimetic values [11]. In contrast, highly transmural infarcts can generate new VTs when covered by the EHT, as shown by Fassina et al. [16] and here. These VTs are suppressed when conductivity is increased.

Overall, our results indicate that the infarct substrate is a primary determinant of VT incidence after remuscularization. For highly transmural SZs, arrhythmicity is reduced when the EHT is implanted laterally to the infarcted region compared to direct implantation over the scar. In contrast, for weakly transmural SZs, VT burden depends on EHT location relative to the infarct, which emphasizes the need for patient-specific digital twins to assess arrhythmic risk and optimize implantation strategies.

### 4.6 Limitations and Future Work

#### Model characteristics

Bishop et al. [50] recently highlighted the need for fine discretizations (*≤* 200 µm) to avoid spurious conduction blocks in slow-conducting tissue and to ensure the reproducibility of arrhythmia induction. In our models, median edge lengths below this threshold were achieved within the EHT and adjacent BZ, and conduction velocities were tuned to OM data, consistent with these recommendations. Although coarser discretizations have been used successfully in prior studies [51–53], our approach increases robustness.

Low-resolution CMRs [14] and segmentation uncertainties were mitigated using advanced algorithms and extensive experimental validation [17, 21]. This supports our finding that LCx pigs displayed more transmural SZs than LAD pigs, likely reflecting both differences in MI induction protocol (proximal LCx versus mid-LAD balloon inflation) and higher out-of-plane LGE-CMR resolution in LAD cases, which allowed thinner BZ layers to be identified [17].

Accurate CS representation is confirmed to be essential for reducing variability in arrhythmia inducibility. Accurate personalization is still limited, as detailed CS anatomy can only be obtained from *ex vivo* histology. Endocardial mapping and non-invasive extracellular potential matching remain the most viable methods for CS modeling validation [17, 34, 43, 54]. Future work should incorporate post-MI region-specific CS conduction delays, as recently shown to be highly arrhythmogenic by Sayers et al. [47].

#### Intrinsic heterogeneities of EHTs

Several intrinsic factors [4, 55] may influence outcomes but are simplified here:

- Multicellular composition: Future models could incorporate stromal cardiac cells with fetal-like phenotypes.
- HiPSC-CM variability: Experimental variability in hiPSC-CMs has motivated the development of multiple computational ionic models, resulting in differences in automaticity and susceptibility to arrhythmia [12, 56, 57]. Recently, Riebel et al. [36] computationally demonstrated that the injection of heterogeneous hiPSC-CMs (ventricular, atrial, nodal) facilitated the generation of post-treatment ectopic activity in a BiV model of LAD MI. Extending our work to incorporate such variability could reveal pig-specific differences in VT burden.
- Cell alignment: hiPSC-CMs often aggregate in islands or exhibit scaffold-driven orientation [7, 58]. Here, a random distribution was assumed for hiPSC-CMs in the EHT, consistent with prior slab and BiV simulations showing a limited arrhythmic impact of alignment [11, 18, 20].

#### Extrinsic factors post-implantation of the EHT

External influences also affect remuscularization outcomes [7, 11, 58, 59].

- Degree of attachment: Incomplete physical engraftment may arise from air bubbles in poorly glued constructs, scar tissue from stitching, immune EHT isolation in xenotransplantation approaches, or shear stress during cardiac contraction-relaxation cycles [7, 11]. Simulations suggest that homogeneous partial engraftment increases the VT risk, whereas the risk is comparable for partial and complete engraftment when the attachment is heterogeneous [11, 16]. Consistently, our previous computational study found only slight electrophysiological variations across degrees of heterogeneous engraftment, provided minimal electrical contact existed [19]. Therefore, here we focused on complete engraftment.
- EHT-induced ectopy: Experimental studies report a high ectopic burden after cellular injection [11, 27, 59]. Computational work indicates that ectopy arises under incomplete engraftment or extreme hiPSC-CM heterogeneity, but not under complete engraftment, and that sVT further requires a bradycardic beat and suppression of CS retrograde conduction [36, 60]. Our results and others [11, 16, 18–20, 48] found no ectopic-driven VT, consistent with the high electrotonic load of the host myocardium suppressing spontaneous activity and the intrinsically slower beating rate of the EHT, ensuring that it is paced by the host myocardium. Recent EHT-related preclinical and clinical studies [27, 61] reporting no arrhythmogenicity support this; however, confirmation of stable EHT–myocardium coupling and high cell survival is still needed. Future studies could explore bradycardic rhythms and heterogeneous grafts to model conditions more favorable to ectopy.
- Transitional phase: We, like others [11, 16], focused on baseline MI and post-EHT implantation endpoints. However, dynamic processes during graft integration, including changes in EHT-myocardium coupling and cell viability, may strongly influence arrhythmogenicity. Future simulations could incorporate time-varying conductivity and hiPSC-CM electrophysiology to capture this transitional phase.

#### Outlook

We developed a realistic EHT-myocardium modeling and simulation framework by integrating an updated hiPSC-CM model with a diverse set of infarct geometries. This approach captures key structural and electrophysiological determinants of arrhythmicity and can be extended as new experimental data emerge. Ultimately, it provides a robust platform to guide case-specific risk assessment and the optimization of post-MI remuscularization strategies.

## 5 Conclusion

We developed a novel computational pipeline to generate porcine BiV digital twins of infarcted hearts before and after EHT remuscularization, aimed at analyzing VT incidence following therapy. Simulated arrhythmic patterns closely matched experimental data. Larger LAD infarcts were linked to higher arrhythmogenicity. CS representation critically influenced post-MI outcomes. Following EHT implantation, VT burden increased in highly transmural scars, particularly when engraftment occurred directly over the infarct; however, it decreased when EHTs were implanted laterally. In less transmural scars, the relationship between VT inducibility and EHT location was highly variable, highlighting digital twins as a tool for personalized risk prediction. Finally, biomimetic EHT conductivity emerged as the key factor in mitigating VT burden after remuscularization therapy.

## Supporting Information

**S1 Table. Results of the arrhythmia inducibility protocol obtained in the G1 simulations.** G1 simulations correspond to the baseline MI-related evaluation. LCx pigs (4-7) are shown at the left table, and LAD pigs (8-12) are shown at the right table. PS: pacing site, NC: no capture, NR: no reentry, nsVT: non-sustained VT, sVT: sustained VT.

**S2 Table. Results of the arrhythmia inducibility protocol obtained in the G2 simulations.** G2 simulations correspond to the assessment of the CS presence. LCx pigs (6-7) are shown at the top of the table, and LAD pigs (11-12) are shown at the bottom of the table. PS: pacing site, NC: no capture, NR: no reentry, nsVT: non-sustained VT, sVT: sustained VT. Note that pacing sites which resulted in NC in the G1 simulations were excluded (-).

**S3 Table. Results of the arrhythmia inducibility protocol obtained in the G3 simulations.** G3 simulations correspond to the assessment of the remuscularization and the location of the EHT implantation. LCx pigs (4-7) are shown at the left table, and LAD pigs (8-12) are shown at the right table. PS: pacing site, MI: baseline pre-EHT, L1: EHT at L1, L2: EHT at L2, NC: no capture, NR: no reentry, nsVT: non-sustained VT, sVT: sustained VT. Note that a low EHT conductivity was used, and pacing sites that resulted in NC in the G1 simulations were excluded (-).

**S4 Table. Results of the arrhythmia inducibility protocol obtained in the G4 simulations.** G4 simulations correspond to the assessment of the EHT conductivity variation when the EHT was implanted in different locations. The left table correspond to the EHT location L1 and the right table correspond to the EHT location L2. LCx pigs (4-7) are shown at the top of the tables, and LAD pigs (8-12) are shown at the bottom of the tables. PS: pacing site, NR: no reentry, nsVT: non-sustained VT, sVT: sustained VT, C10/90: EHT with a conductivity equal to the 10/90% of the value found in the healthy adult-like tissue.

**S5 Table. Results of the characterization of the MI substrate.** In the left table, the volumetric ratios between the MI and the BiV, as well as, between the BZ and the MI are presented. In the right table, the epicardial surface ratio of the SZ with whole MI is presented. LCx pigs (4-7) are depicted in the top of the tables while the LAD pigs (8-12) can be found at the bottom of the tables.

**S1 Video. Example results of G1 simulations.** Experimental (top) and simulated (bottom) reentrant activity in LAD pigs.

**S2 Video. Example results of G2 simulations.** Simulated reentrant activity for pigs 6 (top-left), 7 (top-right), 11 (bottom-left), and 12 (bottom-right) with (left) and without (right) the CS.

**S3 Video. Example results of G3 simulations.** Simulated reentrant activity for pigs 4 (top), 5 (middle), and 8 (bottom), previous the EHT implantation (left, MI) and following the central (middle, L1) and lateral (right, L2) to the infarction EHT engraftment.

**S4 Video. Example results of G4 simulations.** Simulated reentrant activity when the EHT conductivity was set to 10% (left) and 90% (right) of that in healthy tissue. Results for pigs 4 (top) and 6 (bottom), when the EHT was implanted lateral (L2) and central (L1) to the infarction, respectively, are shown.

